# Interannual variability of net ecosystem carbon production and its climatic and biotic mechanism during 2005-2018 in a rain-fed maize ecosystem

**DOI:** 10.1101/2020.08.03.233973

**Authors:** Hui Zhang, Tianhong Zhao, Sidan Lyu, Hang Wu, Yang Yang, Xuefa Wen

## Abstract

The interannual variation (IAV) of net ecosystem carbon production (NEP) plays an important role in understanding the mechanisms of the carbon cycle in the agriculture ecosystem. In this study, the IAV of NEP, which were expressed as annual values and anomalies, and its climatic and biotic controls mechanism, were investigated based on an eddy covariance dataset of rain-fed spring maize during 2005–2018 in the northeast of China. The annual NEP was 270±115 g C m^−2^yr ^−1^. Annual values and anomalies of NEP were positively correlated with that of precipitation (PPT), gross ecosystem production (GEP) and daily maximum NEP (NEP_max_). Annual anomalies of NEP were dominantly and positively controlled by the soil water content (SWC) through GEP and the soil temperature (Ts) through RE. In comparison, annual anomalies of NEP were dominantly and negatively controlled by summer VPD through the NEP_max_, positively adjusted by spring precipitation and the effective accumulative temperature through the beginning date (BDOY) of the affecting carbon uptake period (CUP), and by autumn precipitation and leaf area index through the end date (EDOY) of the affecting CUP. Residues restrained the carbon release at the beginning of the year, and accelerated the carbon release at the end of the year. Our results hightlight that NEP might be more sensitive to the change of water condition (such as PPT, SWC and VPD) induced by the climate changes.

## 1. Introduction

The interannual variation (IAV) of net ecosystem carbon production (NEP) plays an important role in understanding the mechanisms of the carbon cycle, and it focuses on the analysis of annual average values and anomalies [1-4]. Cultivated land occupies approximately 40% of the terrestrial ecosystem and 30% is used for agriculture [5]. In the Northern Hemisphere, croplands had positive contributions to the long term trend of land carbon uptake with 21%, and it played an important role in the IAV of NEP in the terrestrial ecosystem [1]. Agricultural ecosystems may fluctuate among a net carbon source, a sink, or neutral under the influences of climate change [6]. However, the understanding of IAV of NEP will be limited under the directly considering of the climatic and biotic effects, because of the difficulty in partitioning its numerous causes [7]. The changes of vegetation phenology and physiological processes, driven by the variations of climatic and biotic factors, lead to the IAV of NEP [8]. Understanding the controlling mechanism of climatic and biotic factors for the IAV of NEP, which can be expressed as the difference between gross ecosystem productivity (GEP) and ecosystem respiration (RE) [9-13], or the integration of carbon uptake or release peak and corresponding duration [1, 14-16], is crucial for identifying and predicting the carbon cycle.

Generally, it is necessary to explore the causes of variation in GEP and RE for understanding the possible causes of the IAV of NEP under the driving of climatic and biotic factors, because GEP and RE linked over long timescales at the ecosystem level have the different influences of climatic and biotic factors [4, 13, 15, 17]. Based on the dataset of 544 site-years from 59 sites, it was found that the anomalies of NEP had the potential to show more sensitivity to the climatic and biotic factors of driving GEP, compared to RE [4]. Dold et al.[18] indicated that precipitation increased GEP and residues increased RE in three different agro-ecosystems with corn, soybeans and tallgrass prairie of USA from 2006 to 2015. Chu et al. [19] suggested that the low soil moisture accompanied with high salinity inhibited plant biomass, which drived GEP and RE in reclaimed coastal wetlands. Yang et al. [20] found GEP was limited more than RE by drought stress in grasslands of the Loess Plateau, and the reduction in soil moisture result in weaker sensitivity of RE being dependent on soil temperature. However, there are disagreements regarding climatic and biotic influences on GEP and RE because of the different ecosystems, regions, and climate conditions. Additional attention is needed for improving our understanding of the mechanisms on the IAV of NEP [15, 21, 22].

Furthermore, NEP is defined as the integration of the carbon uptake or release peak (plant physiology indicator) and a corresponding duration (phenological parameter) to illustrate the climatic and biotic influences on the IAV of NEP[1, 14-16]. The framework of peak value and duration provides a simple mechanism for understanding the influences of climatic and biotic variables on the IAV of NEP [15, 16]. Fu et al. [14] disentangled the effects of climatic drivers on NEP by decomposing annual NEP into daily maximum NEP (NEP_max_) and a carbon uptake period (CUP), based on a similar GEP partitioning method reported by Xia et al. [23] and Zhou et al. [8]. Based on the atmospheric inversion dataset and the global analysis, it was found that the large IAV of NEP attributed to both the extended CUP and the increasing amplitude of NEP_max_ [1]. Fu et al. [16] found that NEP was predominately determined by NEP_max_ at the global scale, based on the long-term observed NEP from 66 eddy covariance sites and global products derived from FLUXNET observations. In all, NEP_max_ was determined by summer precipitation, and CUP was determined by the spring temperature during the beginning date of CUP (BDOY) and the autumn temperature, radiation and precipitation during the end date of CUP (EDOY) [8, 14].

Meanwhile, the air and soil temperature, soil moisure and residues was the predominant influencing factors of carbon releases [24-26]. Fu et al. [16] also suggested that 10% of the IAV of NEP is derived over the carbon release period. Suyker and Verma [26] showed that carbon release contributed 10–20% and 17–24% of IAV in maize and soybean agroecosystems, respectively, and was mainly controlled by the air temperature and residue biomass. However, the carbon realease period is not always continuous in a calender year, expecially in the agriculture ecosystem, considered as an integral ecosystem [27].

For the IAV of cropland NEP in China, existing studies were mostly focus on winter wheat and summer maize double-cropped croplands in the North Plain, or spring maize in the Loess Plateau [11, 28-30]. In northeast China, the average growth rate of air temperature is 0.35°C/10a, and that of precipitation is –13.3 mm/10a, based on the 1961–2010 ground observations from 91 meteorological stations [31, 32]. However, the rain-fed spring maize is the dominant crop, and it accounts for 57.2% of the planted area and 64.3% of the crop yield in northeast China, and 33.8% of the total maize yield in China [31, 33].

In this study, continuous measurements of carbon flux by eddy covariance were conducted in rain-fed spring maize in northeast China since June 2004. The IAV of NEP and its climatic and biotic controlling mechanism, which were expressed as annual values and anomalies, were investigated during 2005–2018. NEP were partitioned into the differences between GEP and RE, or the integration of carbon uptake or release peak and corresponding duration. Meanwhile, the effective air accumulative temperature and biotic factors were also considered in this study according to the law of accumulative temperature related to crop growth.

## 2. Materials and Methods

### 2.1 Site characteristics and agricultural management

As a part of the China FLUX network, the eddy fluxes of carbon dioxide and water vapor of a typical rain-fed maize agroecosystem were measured at the Jinzhou site (41°08’ N, 121°12’E, and 23.3 m above sea level) in the northeast China. This flux site is operated by Jinzhou Ecology and Agriculture Meteorological Center, Liaoning Meteorological Bureau, and located in a representative temperate monsoon climate area with an mean annual temperature of 9.4 °C and annual precipitation of 565.9 mm, obtained according to the long-term records (1981–2010) of the adjacent weather station [33]. There were two prevailing wind directions, including north-northeast in winter and south-southeast in summer.

The rain-fed maize was generally sown during mid-April to mid-May and harvested during mid-September to early November every year, depended on air temperature and soil moisture condition without irrigation. Maize was cultivated at a density of 48500 plants per ha. The maximum leaf area index (LAI) and canopy height were 4.4 during the tasseling period and 2.8 m during the filling period, respectively. During the off-season from October/November to April/May, the crop residues were left on the bare soil surface. Bulk density of the loamy soil was 1.6 g cm^−3^, pH was 6.4, and soil organic carbon was 23.6 g kg^−1^ [34]. Chemical fertilizer (NH_4_HCO_3_) was only applied to farmlands during the planting day with the concentrations of 1000 kg ha^−1^ [35].

### 2.2 Eddy covariance and auxiliary measurements

#### 2.2.1 Eddy covariance measurements

The eddy covariance (EC) system was equipped with an open path infrared gas analyzer (LI-7500, Licor Biosciences Inc.) and a 3-D sonic anemometer (CSAT, Campbell Scientific Inc.), which were installed at a fixed height of 4 m. The sampling frequency was 10 Hz and all signals were recorded on data loggers (CR5000, Campbell Scientific Inc.) since June 2004. A gas analyzer was periodically calibrated for dealing with the instrument drift. The flux tower was approximately located in the center of the maize field with at least a 380 m radius. The representative area of the flux footprint climatology decreasing with the growth of maize was 0.02–0.34 km^−2^ in the growing season, and 0.39–0.42 km^−2^ in the non-growing season, calculated by the KM footprint model using data from 2005–2018 [36, 37]. The 80% of flux contribution came from the considered field during the growing season (Fig. S1), and 70% was designated to occur during the non-growing season.

#### 2.2.2 Auxiliary meteorological, soil and plant measurements

Auxiliary radiation measurements were conducted using a 4-component net radiometer (CNR-1, Kipp & Zonen) and a quantum sensor of photosynthetically active radiation (PAR) (Li190SB, LI-COR Inc.), positioned over the canopy at the heights of 5 m and 4 m, respectively. Precipitation (PPT) was monitored by a tipping bucket (52202, RM Young Inc). Air temperature (Ta) and relative humidity were measured by an HMP45C (Vaisala, Campbell Scientific Inc.) at the heights of 4 m and 6 m. Wind speed and direction were measured by a wind sensor (034B,MetOne Inc.) at the heights of 6 m. Soil volumetric water content (SWC) at 10 cm depth was measured by Eazy-AG50 (Sentek Inc.) and soil temperature (Ts) at the depths of 5, 10, 15, 20, 40 and 80 cm were obtained by HFT01SC (Campbell Scientific Inc.). All of the meteorological variables were collected using a datalogger (CR23X, Campbell Scientific Inc.).

The LAI was determined according to the corn observation specifications of the China Meteorological Administration. In summary, 5 maize plants at each growth stage were randomly sampled. The lengths and widths of every leaf were manually measured for each sample. The leaf areas of the samples were averaged with a maize coefficient 0.7 [38]. Then, the average leaf area was combined with the plant density to obtain the LAI during the every growth stage. The daily LAI was calculated by the logistic equation interpolation [31], and then the monthly LAI during the maize growth stage period was obtained. The monthly and annual LAI were obtained by the relationship between monthly LAI and monthly NDVI from MODIS during the maize growth stage period.

The residues were defined as the carbon content of leaves and stems left on the bare soil surface during the off-season, and the roots were ignored. 40 maize plants after maturity were randomly sampled and divided into grains, stems, and leaves after 30 days of drying. The corresponding weights per unit area were assessed by combining with the plant densities. The carbon content of stems and leaves was determined by their weights multiplied by 0.4 [28]. Organic carbon in the soil was determined by a wet oxidation method [39].

### 2.3 Flux data processing and calculations

#### 2.3.1 Data processing and quality control

Net exchange carbon production (NEP, g C m^−2^ s^−1^) was measured using the EC technique [9, 40, 41].

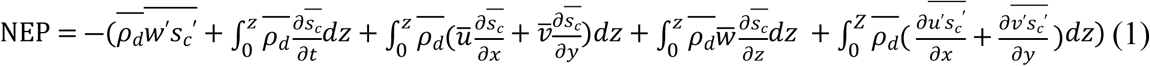

where the terms on the right side of the equation are eddy flux (Term I), storage (Term II), horizontal advection (Term III), vertical advection (Term IV), and horizontal flux divergence (Term V) [9, 41], respectively. Terms III - V are ignored because of the flat and homogeneity assumption [40]. In agriculture ecosystem, Term II was small and therefore neglected in this study.

For the 10 Hz raw data, statistical tests was used to remove spikes [42]. Two-dimension coordinate rotation [43] was conducted to achieve the consistency of coordinates between the instrument and the flow. A Webb, Pearman & Leuning correction was applied to remove the effect of fluctuation in the air density on the fluxes [44]. The low and high frequency spectral corrections were also calculated [45]. The 0-1-2 flags system developed by Mauder and Foken [46] based on the steady state and turbulent condition, was used to classify the flux quality. The data classified with flag 2 was removed. When the friction velocity (u*) was less than the u*-threshold, the flux data was also rejected to avoid possible underestimation of the flux during stable conditions at night. The u*-threshold was determined by the method of Reichstein et al. [12]. These processes were operated in EddyPro software (version6.2.2, LI-COR, Lincoln, NE). The amount of effective data of 2005-2018 after the quality control was shown in Table S1.

#### 2.3.2 Gap filling and flux partitioning

To acquire a continuous dataset and estimate the annual fluxes, marginal distribution sampling was used to fill data-gaps [12]. The NEP is partitioned into GEP and ecosystem respiration (RE). The night NEE is described as RE, because there is minimal photosynthesis at night. The RE is calculated according to the Lloyd and Taylor [47] equation:

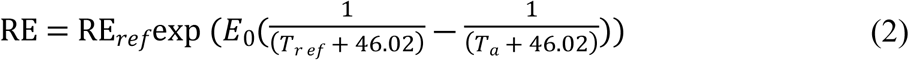

where T_*ref*_ is the reference temperature (10°C), E_0_ is the temperature sensitivity coefficient (°C) and RE_*ref*_ (g C m^−2^ s^−1^) is the reference respiration at T_*ref*_. RE_*ref*_ and E_0_ is estimated by the regression analysis every 15 days [12] and RE of daytime were calculated. Subsequently, GEP is calculated as the sum of NEP and RE. The above processes were conducted using Tovi software (version2.7.2, LI-COR, Lincoln, NE).

### 2.4 Definitions of the carbon uptake or release peak and corresponding duration

#### 2.4.1 Definitions for α, NEP_max_, CUP, and β, NEP_min_ and CRP

For each year, NEP can be decomposed into a net CUP and two net carbon release periods (CRP) at the beginning of the year (CRP__begin_) and the end of the year (CRP__end_, Fig. 1). The CUP was defined as the days between the BDOY and EDOY, and BDOY and EDOY represented the first and last day of positive NEP in a year[1, 8, 14, 16, 23], respectively. The maximum daily NEP (NEP_max_) was defined as the maximum value of daily NEP during CUP, and the minimum daily NEP (NEP_min_begin_ or NEP_min_end_) was defined as the minimum value of daily NEP during the release period (CRP__begin_ or CRP__end_). α was defined as the ratio of actual carbon uptake and hypothetical maximum carbon uptake, and β was defined as the ratio of actual carbon release and hypothetical maximum carbon release [14]. Note that the 10-day moving average was applied to determine the BDOY, EDOY, NEP_max_, NEP_min_begin_ and NEP_min_end_. Annual NEP can be described as a function of the six indicators including α, NEP_max_, CUP, β, NEP_min_, and CRP. The relative contributions of α, NEP_max_, CUP, β and NEP_min_ to NEP anomalies were determined by the specific calculation method reported by Fu et al. [16].

**Fig. 1.**
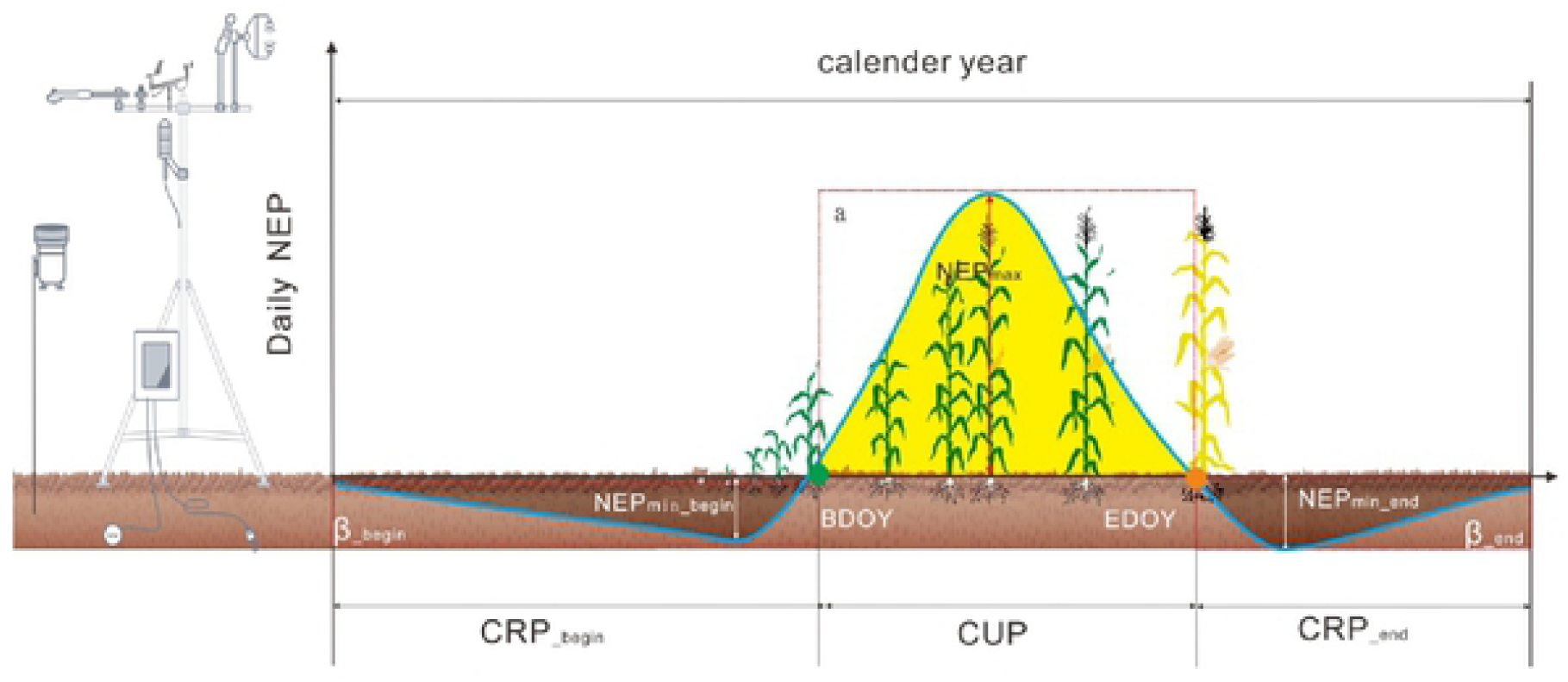
Conceptual figure of the ratio of actual carbon uptake and hypothetical maximum carbon uptake (α), maximum daily NEP (NEP_max_), carbon uptake period (CUP), and the ratio of actual carbon release and hypothetical maximum carbon release (β), minimum daily NEP (NEP_min_) and carbon release period (CRP) at the beginning and end of the year in determining the changes in annual net ecosystem CO_2_ exchange.

#### 2.4.2 Definitions for climatic and biotic variables

The whole year was divided into winter (December–February), spring (March–May), summer (June–August), and autumn (September–November). The seasonal and annual values of carbon fluxes (NEP, GEP, and RE), climatic variables (PAR, PPT, VPD and Ta), soil variables (Ts at 5cm and SWC) and biotic variables (LAI and Resi) were calculated to estimate the seasonal climatic and biotic controls in response to the variability of NEP. According to the law of accumulative temperature, crops need a range of temperature accumulation to accomplish a stage of development. Thus, the effective air accumulative temperature (Ta__ef_) was analyzed, which was greater than 10°C.

### 2.5 Statistical analyses

If variables had a significant linear regression with time, the annual anomalies were defined as the differences between annual values and the values from the linear regression with time, based on the procedure of linear detrending. Otherwise, the annual anomalies were considered as the differences between the annual values related to long-term average results.

Linear correlation was used to evaluate the influences of climatic and biotic variables on NEP (GEP and RE or carbon uptake/release peak value and corresponding duration). Redundancy analysis (RDA) was applied to estimate the contribution of climatic and biotic variables to the annual anomalies of NEP. Structural equation modeling (SEM) was employed to investigate the relationships between climatic and biotic variables, the annual anomalies of NEP, GEP and RE, and the uptake and release peak value and corresponding duration. Linear and stepwise regression were performed by SPSS 18.0 for Windows Software (Version 18.0, SPSS Inc., Chicago, IL, USA). RDA analyses were applied in R version 3.6.2. SEM analyses were applied by AMOS 22.0 (Amos Development Corporation, Chicago, IL, USA).

## 3. Results

### 3.1 The IAV of NEP and its climatic and biotic direct controls

The annual values of carbon fluxes (NEP, GEP, and RE, Fig. 2) and climatic (PAR, PPT, VPD and Ta), soil (Ts and SWC) and plant (LAI and Resi) variables (Table 1), and their annual anomalies were estimated. The annual average NEP was 270±115 g C m^−2^yr ^−1^ (Table 1), and there was no significant increasing or decreasing trend, although the great interannual variations were observed. Annual values and anomalies of NEP were positively correlated with PPT, and annual anomalies of NEP were also negatively correlated with PAR and VPD (Table S2). In addition, annual PAR and VPD had the significant increasing trends (Table 1).

**Table 1.**
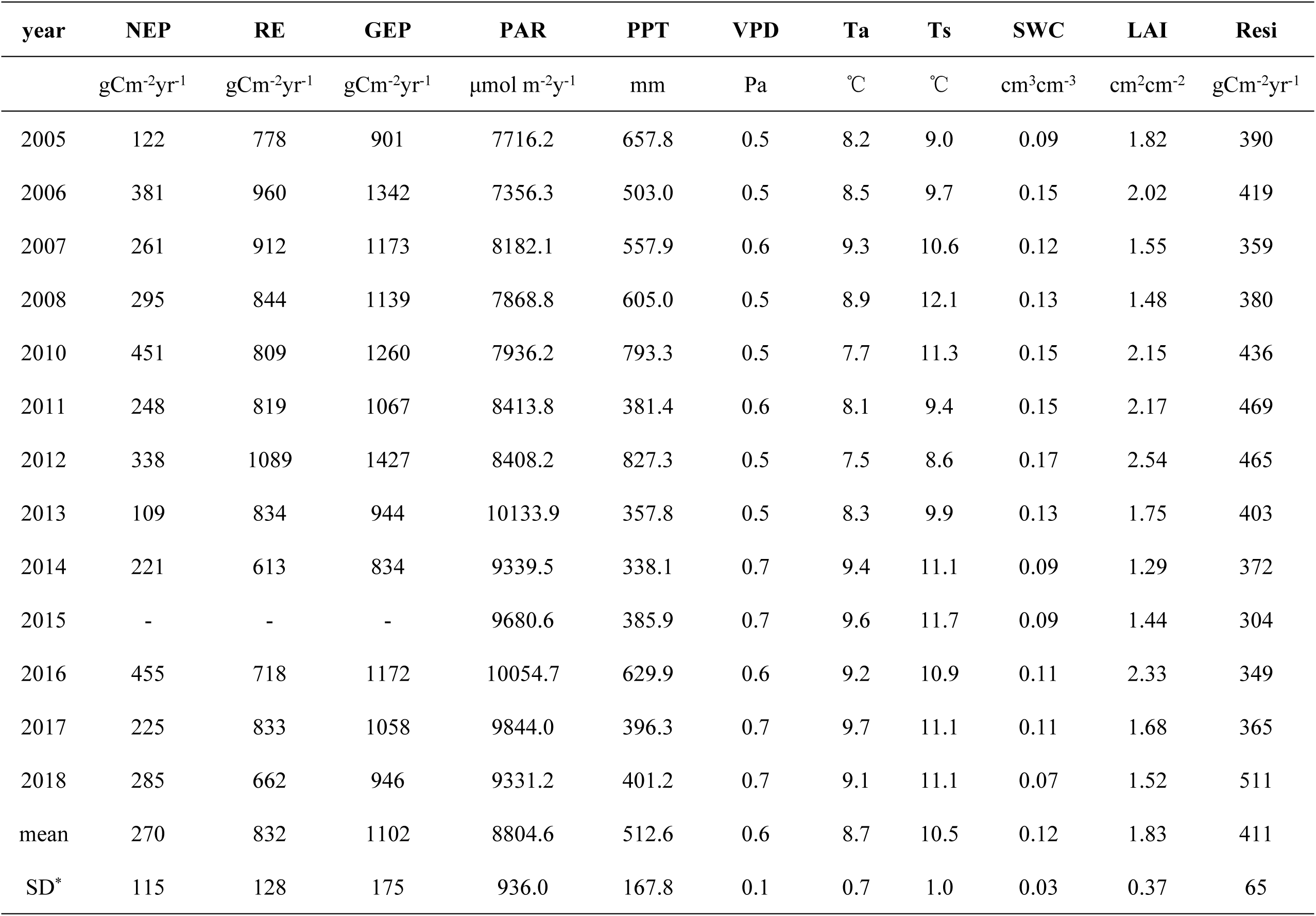
Annual values of net ecosystem production (NEP), ecosystem respiration (RE), gross ecosystem production (GEP), photo synthetically active radiation (PAR), precipitation (PPT), vapor pressure deficit (VPD), air and soil temperature (Ta, Ts), soil water content (SWC), leaf area index (LAI) and residue (Resi). SD means standard deviation.

| year | NEP | RE | GEP | PAR | PPT | VPD | Ta | Ts | SWC | LAI | Resi |
| --- | --- | --- | --- | --- | --- | --- | --- | --- | --- | --- | --- |
|  | gCm<sup>-2</sup>yr<sup>-1</sup> | gCm<sup>-2</sup>yr<sup>-1</sup> | gCm<sup>-2</sup>yr<sup>-1</sup> | μmol m<sup>-2</sup>y<sup>-1</sup> | mm | Pa | °C | °C | cm<sup>3</sup>cm<sup>-3</sup> | cm<sup>2</sup>cm<sup>-2</sup> | gCm<sup>-2</sup>yr<sup>-1</sup> |
| 2005 | 122 | 778 | 901 | 7716.2 | 657.8 | 0.5 | 8.2 | 9.0 | 0.09 | 1.82 | 390 |
| 2006 | 381 | 960 | 1342 | 7356.3 | 503.0 | 0.5 | 8.5 | 9.7 | 0.15 | 2.02 | 419 |
| 2007 | 261 | 912 | 1173 | 8182.1 | 557.9 | 0.6 | 9.3 | 10.6 | 0.12 | 1.55 | 359 |
| 2008 | 295 | 844 | 1139 | 7868.8 | 605.0 | 0.5 | 8.9 | 12.1 | 0.13 | 1.48 | 380 |
| 2010 | 451 | 809 | 1260 | 7936.2 | 793.3 | 0.5 | 7.7 | 11.3 | 0.15 | 2.15 | 436 |
| 2011 | 248 | 819 | 1067 | 8413.8 | 381.4 | 0.6 | 8.1 | 9.4 | 0.15 | 2.17 | 469 |
| 2012 | 338 | 1089 | 1427 | 8408.2 | 827.3 | 0.5 | 7.5 | 8.6 | 0.17 | 2.54 | 465 |
| 2013 | 109 | 834 | 944 | 10133.9 | 357.8 | 0.5 | 8.3 | 9.9 | 0.13 | 1.75 | 403 |
| 2014 | 221 | 613 | 834 | 9339.5 | 338.1 | 0.7 | 9.4 | 11.1 | 0.09 | 1.29 | 372 |
| 2015 | - | - | - | 9680.6 | 385.9 | 0.7 | 9.6 | 11.7 | 0.09 | 1.44 | 304 |
| 2016 | 455 | 718 | 1172 | 10054.7 | 629.9 | 0.6 | 9.2 | 10.9 | 0.11 | 2.33 | 349 |
| 2017 | 225 | 833 | 1058 | 9844.0 | 396.3 | 0.7 | 9.7 | 11.1 | 0.11 | 1.68 | 365 |
| 2018 | 285 | 662 | 946 | 9331.2 | 401.2 | 0.7 | 9.1 | 11.1 | 0.07 | 1.52 | 511 |
| mean | 270 | 832 | 1102 | 8804.6 | 512.6 | 0.6 | 8.7 | 10.5 | 0.12 | 1.83 | 411 |
| SD* | 115 | 128 | 175 | 936.0 | 167.8 | 0.1 | 0.7 | 1.0 | 0.03 | 0.37 | 65 |

**Fig. 2.**
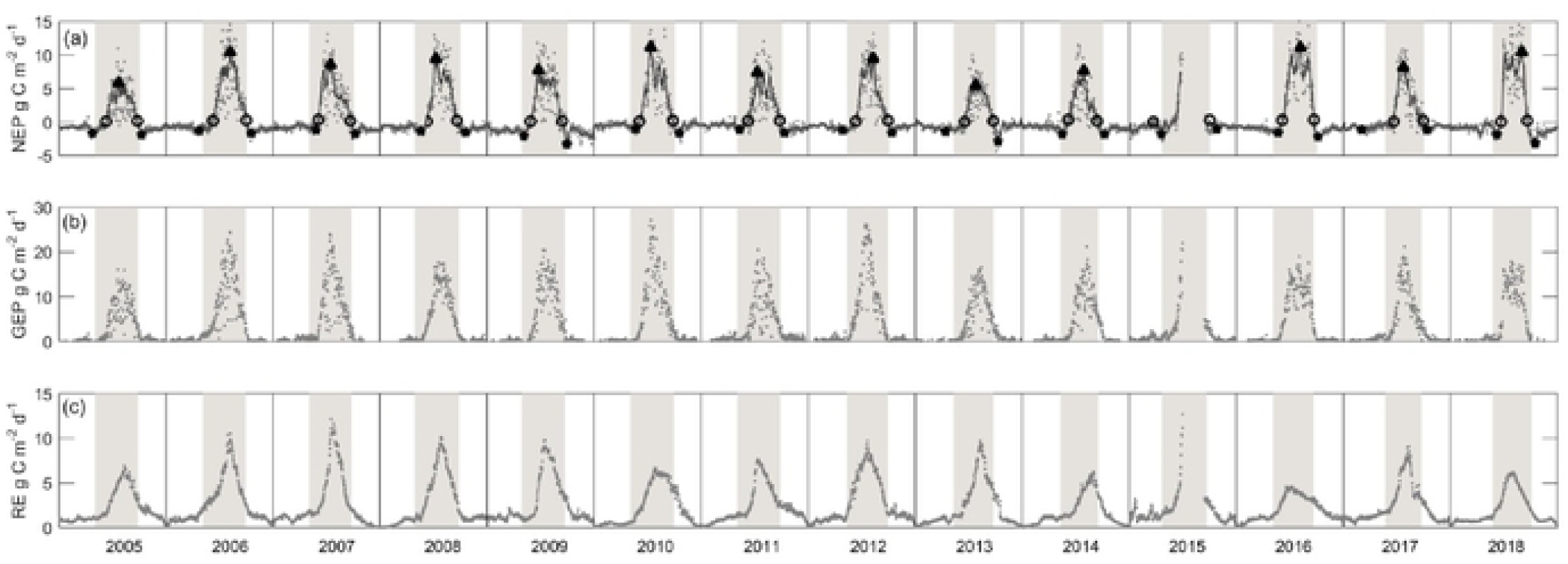
Seasonal and interannual dynamics of daily integrated (a) net ecosystem production (NEP), (b) gross ecosystem production (GEP) and (c) ecosystem respiration (RE) from 2005 to 2018. Gray lines indicate 10-day moving average NEP, black triangle indicates maximum daily net ecosystem production (NEP_max_), black circles indicate the beginning and ending date of net carbon uptake (BDOY and EDOY), black points indicate minimum daily NEP (NEP_min_).

RDA analysis showed that 78.9% of annual anomalies of NEP were explained by the interactions of climate factors (PPT, VPD and Ta), soil factors (SWC) and biotic factors (LAI) (P=0.032, Fig. S2). The contributions of climate, soil, and plant factors for the annual anomalies of NEP were 53.9%, 26.9% and 21.7%, respectively. Furthermore, according to SEM results, Annual anomalies of NEP were negatively dominated by VPD, and VPD was negatively adjusted by PPT and positively adjusted by PAR (Fig. S3).

### 3.2 Climatic and biotic controls of the IAV of NEP through GEP and RE

Annual values and anomalies of NEP were positively and significantly correlated with GEP but not with RE (Table S3). This result indicated that the climatic and biotic factors controlling GEP were easier to lead to IAV of NEP, compared with the factors controlling RE. In addition, annual values and anomalies of GEP were positively correlated with PPT (R^2^ = 0.43, P < 0.01), SWC (R^2^ = 0.59, P < 0.01) and LAI (R^2^ = 0.47, P < 0.01), and annual anomalies of GEP were also negatively correlated with VPD (R^2^ = 0.33, P < 0.05) (Table S4). However, the annual values and anomalies of RE were negatively correlated with Ta and Ts, and positively correlated with SWC (Table S5).

The SEM result showed that climatic and biotic controls through GEP and RE explained 88% of annual anomalies of NEP (Fig. 3). Note that GEP and RE directly and bi-directionally affected annual anomalies of NEP. SWC was the main controlling factor for GEP, while Ts was the main controlling factor for RE. Annual anomalies of NEP were dominantly and positively controlled by SWC through GEP and by Ts through RE. Note that SWC was positively controlled by LAI for GEP, while Ts was negatively controlled by LAI for RE. For both GEP and RE, LAI were negatively controlled by VPD, while VPD were negatively controlled by PPT.

**Fig. 3.**
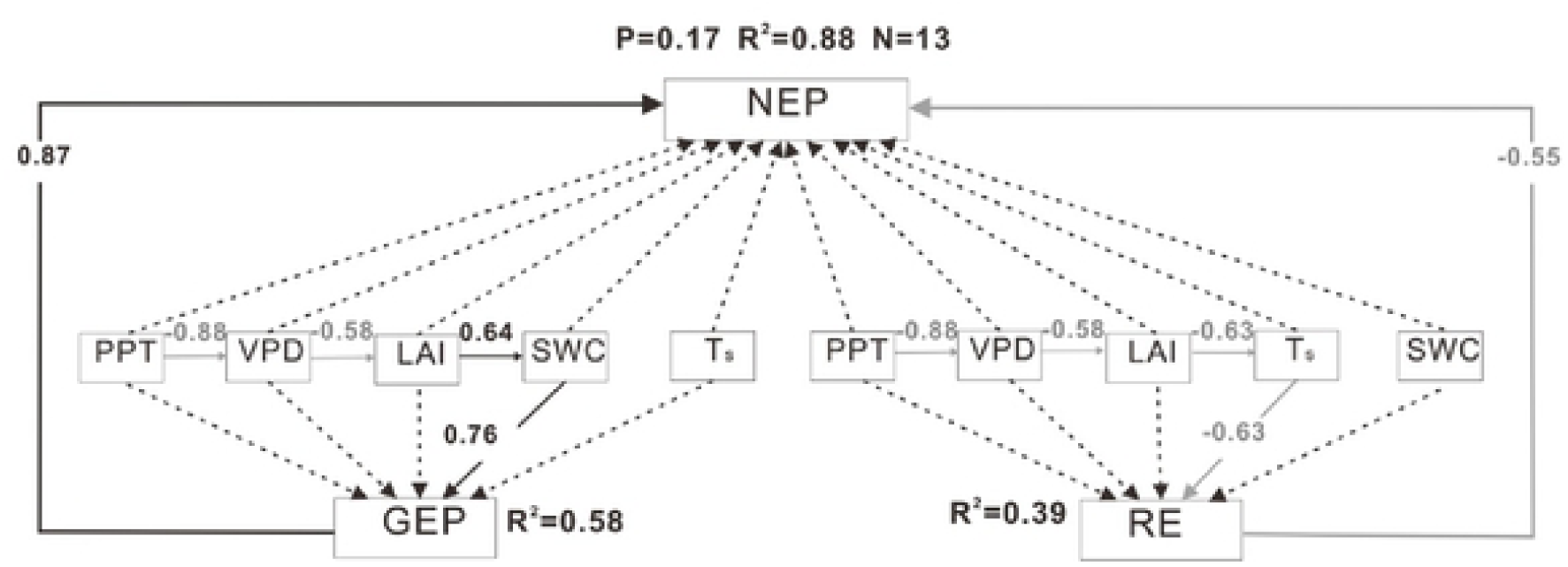
The structure equation modeling results of the relationship among annual anomalies of climatic and biotic variables, gross ecosystem productivity (GEP), ecosystem respiration (RE), and net ecosystem production (NEP). Black arrows indicate significant positive relationships while gray arrows indicate significant negative relationships (P < 0.05). Black dashed arrows indicate insignificant relationships (P > 0.05). Numbers adjacent to arrows are path coefficients and indicative of the effect size of the relationship. The proportion of variance explained (R^2^) appears alongside every response variable in the model.

### 3.3 Climatic and biotic controls of the IAV of NEP through the integration of peak value and duration

#### 3.2.1 The influences of CUP and NEP_max_ on the IAV of NEP during the whole year and the uptake period

Annual values (Table 2) and anomalies of CUP, BDOY, EDOY, NEP_max_, NEP_min_begin_ and NEP_min_end_ were calculated. Annual values and anomalies of NEP were positively correlated with NEP_max_, while annual anomalies of NEP were positively correlated with CUP and negatively with CRP at the beginning of the year (Table 3). Notably, there was a significant decreasing trend in CUP (R^2^ = 0.48, P <0.01) with a significant increasing trend of BDOY (R^2^ = 0.33, P <0.05), except in 2015 (Table 2).

**Table 2.**
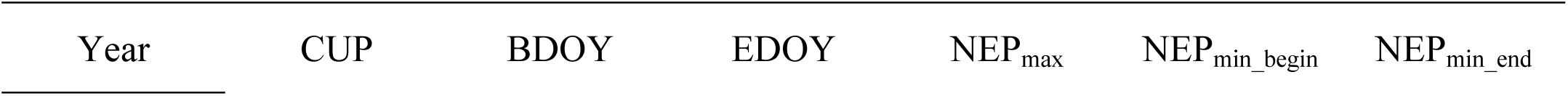
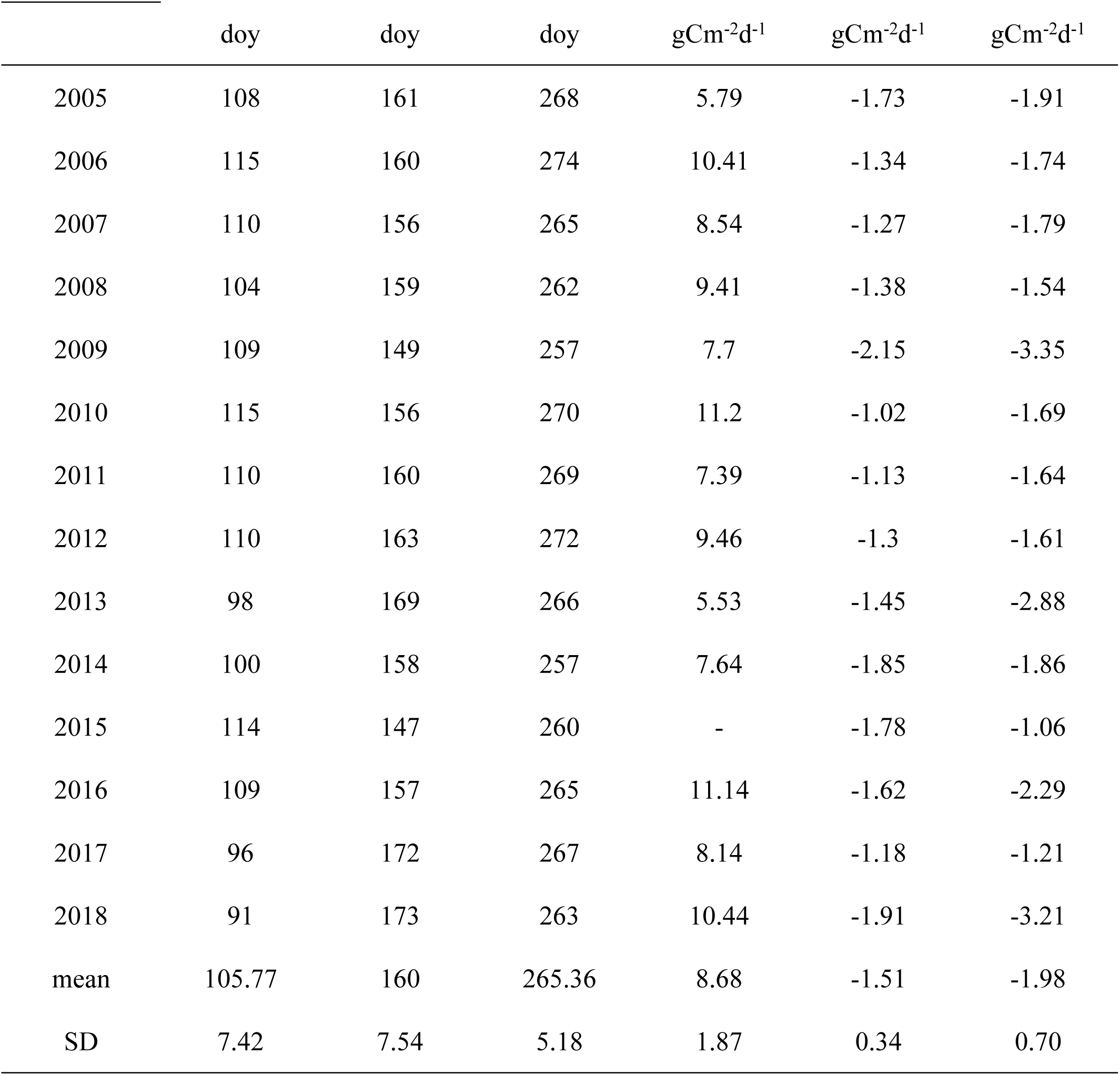
Annual values of carbon uptake period (CUP), the beginning and ending date of net carbon uptake (BDOY and EDOY), maximum daily net ecosystem production (NEP_max_), and minimum daily NEP (NEP_min_begin_ and NEP_min_end_). SD means standard deviation.

|  | doy | doy | doy | gCm<sup>-2</sup>d<sup>-1</sup> | gCm<sup>-2</sup>d<sup>-1</sup> | gCm<sup>-2</sup>d<sup>-1</sup> |
| --- | --- | --- | --- | --- | --- | --- |
| 2005 | 108 | 161 | 268 | 5.79 | -1.73 | -1.91 |
| 2006 | 115 | 160 | 274 | 10.41 | -1.34 | -1.74 |
| 2007 | 110 | 156 | 265 | 8.54 | -1.27 | -1.79 |
| 2008 | 104 | 159 | 262 | 9.41 | -1.38 | -1.54 |
| 2009 | 109 | 149 | 257 | 7.7 | -2.15 | -3.35 |
| 2010 | 115 | 156 | 270 | 11.2 | -1.02 | -1.69 |
| 2011 | 110 | 160 | 269 | 7.39 | -1.13 | -1.64 |
| 2012 | 110 | 163 | 272 | 9.46 | -1.3 | -1.61 |
| 2013 | 98 | 169 | 266 | 5.53 | -1.45 | -2.88 |
| 2014 | 100 | 158 | 257 | 7.64 | -1.85 | -1.86 |
| 2015 | 114 | 147 | 260 | - | -1.78 | -1.06 |
| 2016 | 109 | 157 | 265 | 11.14 | -1.62 | -2.29 |
| 2017 | 96 | 172 | 267 | 8.14 | -1.18 | -1.21 |
| 2018 | 91 | 173 | 263 | 10.44 | -1.91 | -3.21 |
| mean | 105.77 | 160 | 265.36 | 8.68 | -1.51 | -1.98 |
| SD | 7.42 | 7.54 | 5.18 | 1.87 | 0.34 | 0.70 |

**Table 3.** Linear regressions between annual values and anomalies of net ecosystem production (NEP), and that of the carbon uptake and release peak value (NEP_max_, NEP_min_begin_ and NEP_min_end_) and corresponding duration (CUP, CRP__begin_ and CRP__end_). ^**^ and ^*^ represent a significant relationship at p = 0.01, and 0.05 levels, respectively.

| Variables (y) |  | Variables (x) | Linear regressions | R<sup>2</sup> | P |
| --- | --- | --- | --- | --- | --- |
| annual values of<br>NEP | uptake | CUP | y =6.59x - 426.85 | 0.18 | 0.149 |
|  |  | NEP<sub>max</sub> | y =56.90x - 223.34 | 0.85 | **0.000\*\*** |
|  | release | CRP<sub>begin</sub> | y =-2.28x + 636.61 | 0.19 | 0.661 |
|  |  | CRP<sub>end</sub> | y =-8.84x + 1149.6 | 0.16 | 0.183 |
| | | NEP<sub>min_begin</sub> | $y = 149.71x + 492.74$ | 0.2 | 0.129 |
| | | NEP<sub>min_end</sub> | $y = 60.586x + 394.76$ | 0.13 | 0.236 |
| annual anomalies<br>of NEP | uptake | CUP<sub>aomalies</sub> | $y = 15.46x + 0.57$ | 0.51 | **0.006\*\*** |
| | | NEP<sub>max_aomalies</sub> | $y = 56.90x + 2E-14$ | 0.85 | **0.000\*\*** |
| | release | CRP<sub>begin_aomalies</sub> | $y = -13.47x + 23.23$ | 0.36 | **0.040\*** |
| | | CRP<sub>end_aomalies</sub> | $y = -8.84x - 6.02$ | 0.16 | 0.183 |
| | | NEP<sub>min_begin_aomalies</sub> | $y = 149.71x - 13.83$ | 0.2 | 0.129 |
| | | NEP<sub>min_end_aomalies</sub> | $y = 60.59x + 1.28$ | 0.13 | 0.236 |

The SEM analysis showed that 89% of annual anomalies of NEP were explained by climatic and biotic controls through CUP and NEP_max_ during the whole year (Fig. 4), and 91% during the carbon uptake period (Fig. S4). Note that CUP and NEP_max_ directly affected annual anomalies of NEP during the whole year and during the carbon uptake period. Annual anomalies of NEP were dominantly and negatively controlled by summer VPD through NEP_max_. Meanwhile, annual anomalies of NEP were also positively controlled by spring precipitation and the effective accumulative temperature through BDOY and by autumn precipitation and LAI through the EDOY affecting CUP.

**Fig. 4.**
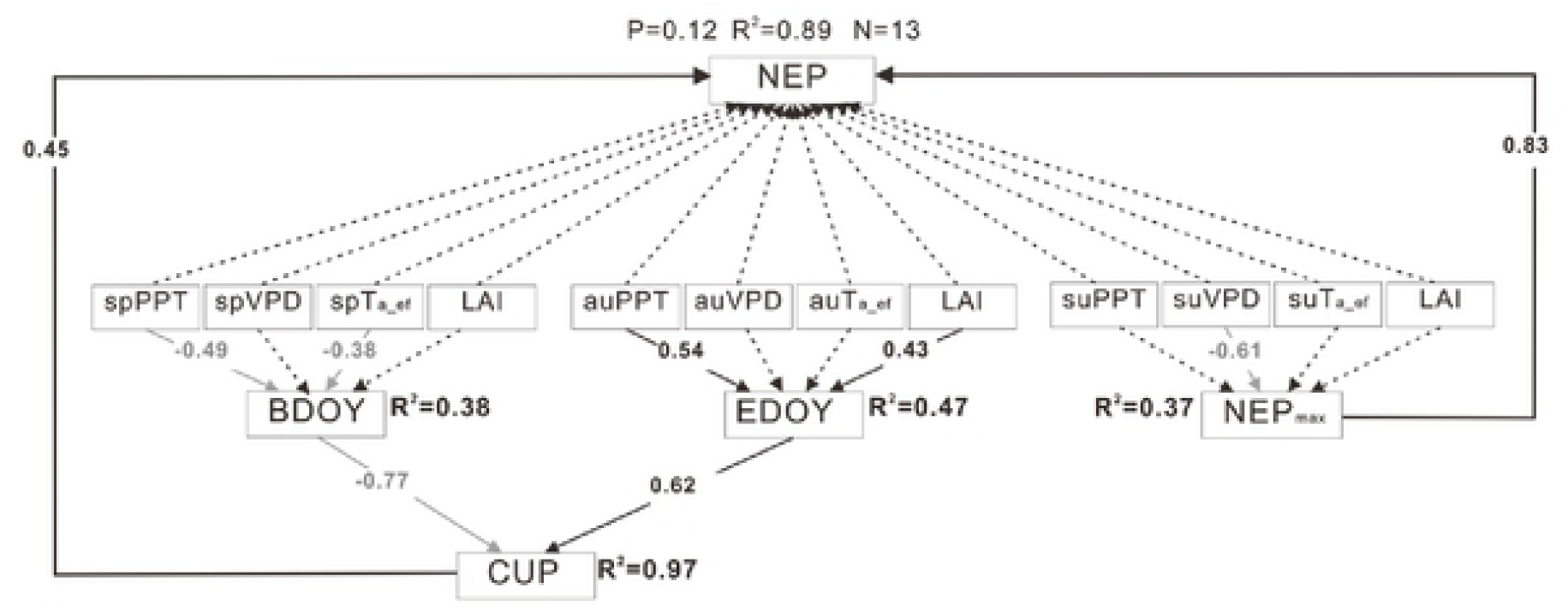
The structure equation modeling results of the relationship among annual anomalies of climatic and biotic variables, carbon uptake period (CUP) and maximum daily net ecosystem production (NEP_max_) and net ecosystem production (NEP) in a single year. Black arrows indicate significant positive relationships while gray arrows indicate significant negative relationships (P < 0.05). Black dashed arrows indicate insignificant relationships (P > 0.05). Numbers adjacent to arrows are path coefficients and indicative of the effect size of the relationship. The proportion of variance explained (R^2^) appears alongside every response variable in the model. sp, su, and au indicate spring, summer, and autumn, respectively

Note that CUP was negatively controlled by BDOY and positively controlled by EDOY. BDOY was negatively correlated with spring precipitation and the effective accumulative temperature. EDOY was positively correlated with LAI and autumn precipitation. NEP_max_ had a negative correlation with summer VPD. The relative contributions of CUP, NEP_max_ and α for annual anomalies of NEP during the uptake period were 9.7%, 84.1% and –3.1% (Table 4), respectively.

**Table 4.**
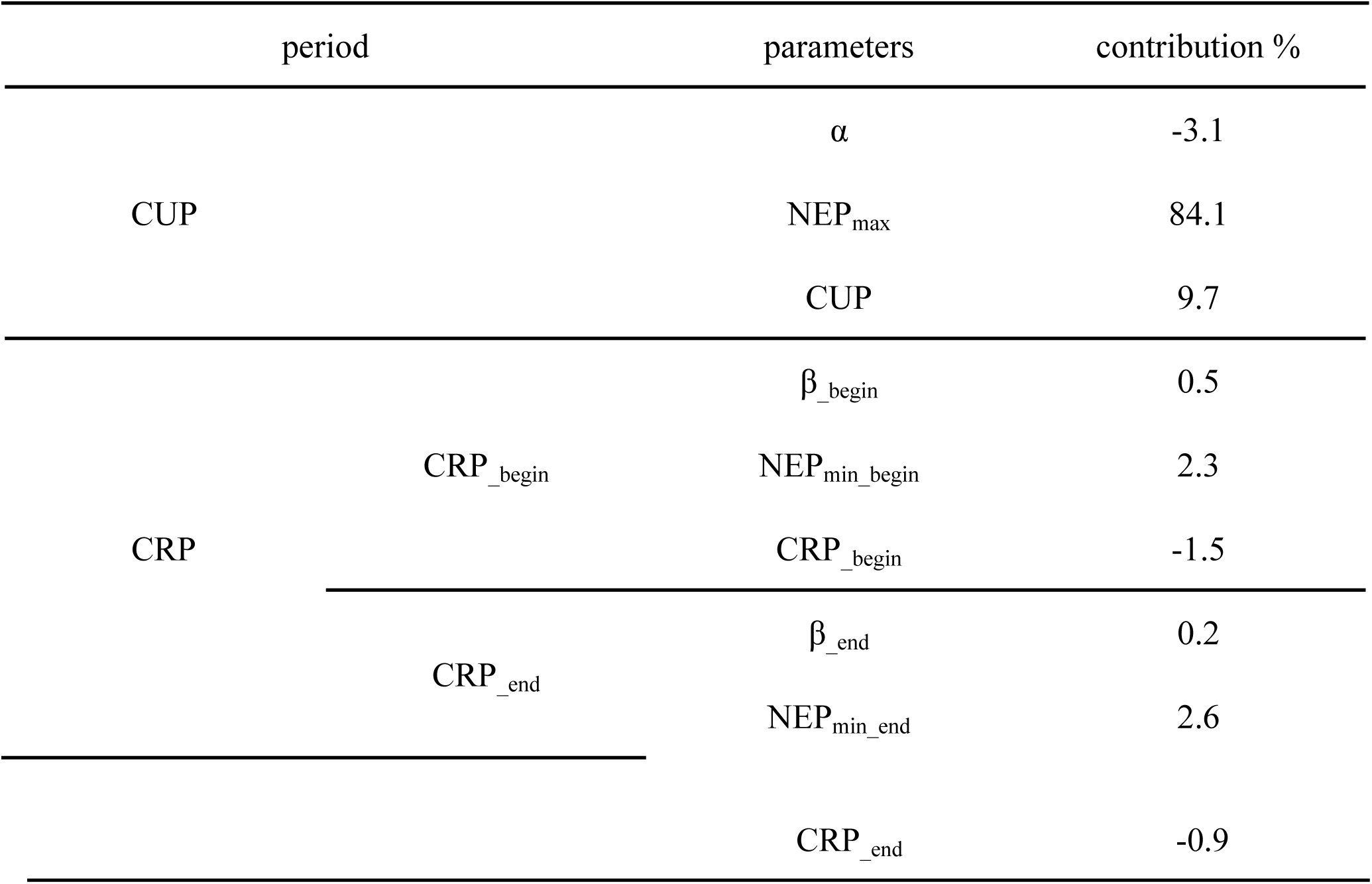
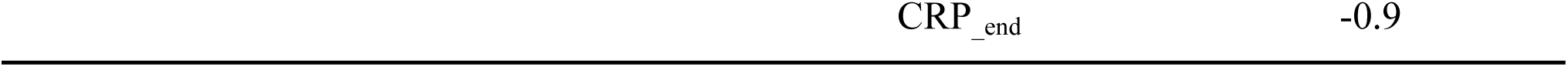
Relative contributions of the ratio of actual carbon uptake and hypothetical maximum carbon uptake (*α*), the maximum daily net ecosystem production (NEP_max_), the net carbon uptake period (CUP), the ratio of actual carbon release and hypothetical maximum carbon release (*β*), the minimum daily NEP (NEP_min_) and carbon release periods (CRP) to annual anomalies of net ecosystem production (NEP)

#### 3.3.2 The influences of CRP and NEP_min_ on the IAV of NEP during release period

The relative contributions of CRP, NEP_min_ and β at the beginning of the year for annual anomalies of NEP were 0.5%, 2.3% and –1.5%, and at the end of the year were 0.2%, 2.6% and –0.9% (Table 4), respectively. 68% of annual anomalies of the carbon release amount at the beginning of the year were explained by climatic and biotic controls through the contemporary CRP and NEP_min_ (Fig. 5a). Annual anomalies of the carbon release amount at the beginning of the year were dominated by NEP_min_ at the beginning of the year, which was restricted by the residues of the previous year.

**Fig. 5.**
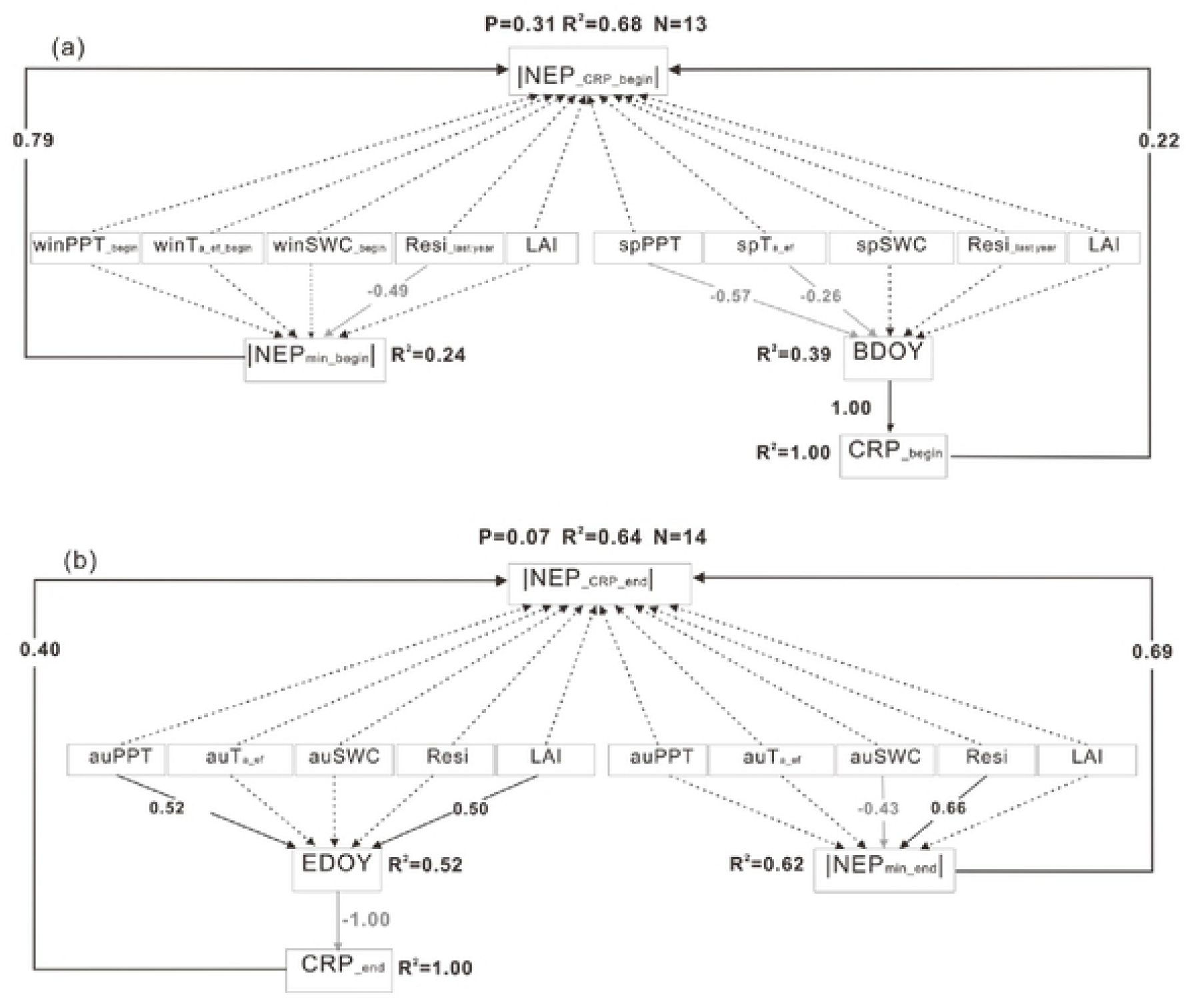
The structure equation modeling results of the relationship among annual anomalies of climatic and biotic variables, carbon release periods (CRP), minimum daily net ecosystem production (NEP_min_) and net ecosystem production (NEP) during release period. (a) represents CRP at the beginning of the year and (b) represents CRP at the end of the year. Black arrows indicate significant positive relationships while gray arrows indicate significant negative relationships (P <0.05). Black dashed arrows indicate insignificant relationships (P > 0.05). Numbers adjacent to arrows are path coefficients and indicative of the effect size of the relationship. The proportion of variance explained (R^2^) appears alongside every response variable in the model. sp, au, and win indicate spring, autumn, and winter, respectively. Resi__last year_ indicates the residues of last year; Resi indicates the residues of current year.

In comparison, 64% of annual anomalies of carbon release amount at the end of the year were explained by climatic and biotic controls through the contemporary CRP and NEP_min_ (Fig. 5b). Annual anomalies of the carbon release amount at the end of the year were dominated by NEP _min_ at the end of the year, which was increased by the residues of the current year and restricted by SWC. Similarly, CRP at the beginning and end of the year were positively correlated with the corresponding amount of the carbon release. The CRP at the beginning of the year was positively correlated with the BDOY, and at the end of the year was negatively correlated with the EDOY.

## 4. Discussions

### 4.1 Partitioning the NEP into the difference between GEP and RE

NEP is the difference between GEP and RE, and the two processes with different physical and biological controls are linked over long timescales at the ecosystem level [9-13]. Partitioning of NEP into GEP and RE is needed for better understanding the causes of IAV of NEP [10]. GEP, as a direct indicator of carbon uptake during ecosystem photosynthesis, is controlled by canopy development, nutrient status, and light, temperature and water conditions [2, 3, 25, 48]. RE, as a direct indicator of carbon loss during ecosystem respiration, is controlled by temperature, soil moisture, nutrient availability, stocks of living and dead biomass, and ecosystem productivity [11, 26, 49]. It is necessary to explore the causes of variation in GEP and RE rather than directly correlate NEP with meteorological or abiotic drivers [13, 26, 50].

GEP is a stronger driver of IAV in NEP compared to RE, and it indicates that annural anomalies of NEP are more sensitive to factors driving GEP compared to those of RE [4, 17, 26, 51]. Baldocchi and Penuelas [21] showed that RE significantly increased with the increasing of GEP, and the slope was 0.82 at the regional scale, and it indicated that 82% assimilated carbon was lost by RE based on the 1163 site-year EC datasets of 155 sites. Luyssaert et al. [52] pointed that the ratio of RE to GEP ranged between 0.76 and 0.97 based on the 1374 site-years EC datasets of 513 sites. Furthermore, the carbon fluxes of 20 sites derived from the published literature on different farmlands at the regional scale (Fig. 6), showed that NEP had a significant positive correlation with GEP, but not with RE, and the ratio of RE and GEP was 0.78 (Table S6). In this study, annual values and anomalies of NEP had the significant positive correlation with GEP, but not with RE. Meanwhile, there was a significant positive correlation between RE and GEP (Table S3) with a ratio of 0.75.

**Fig. 6.**
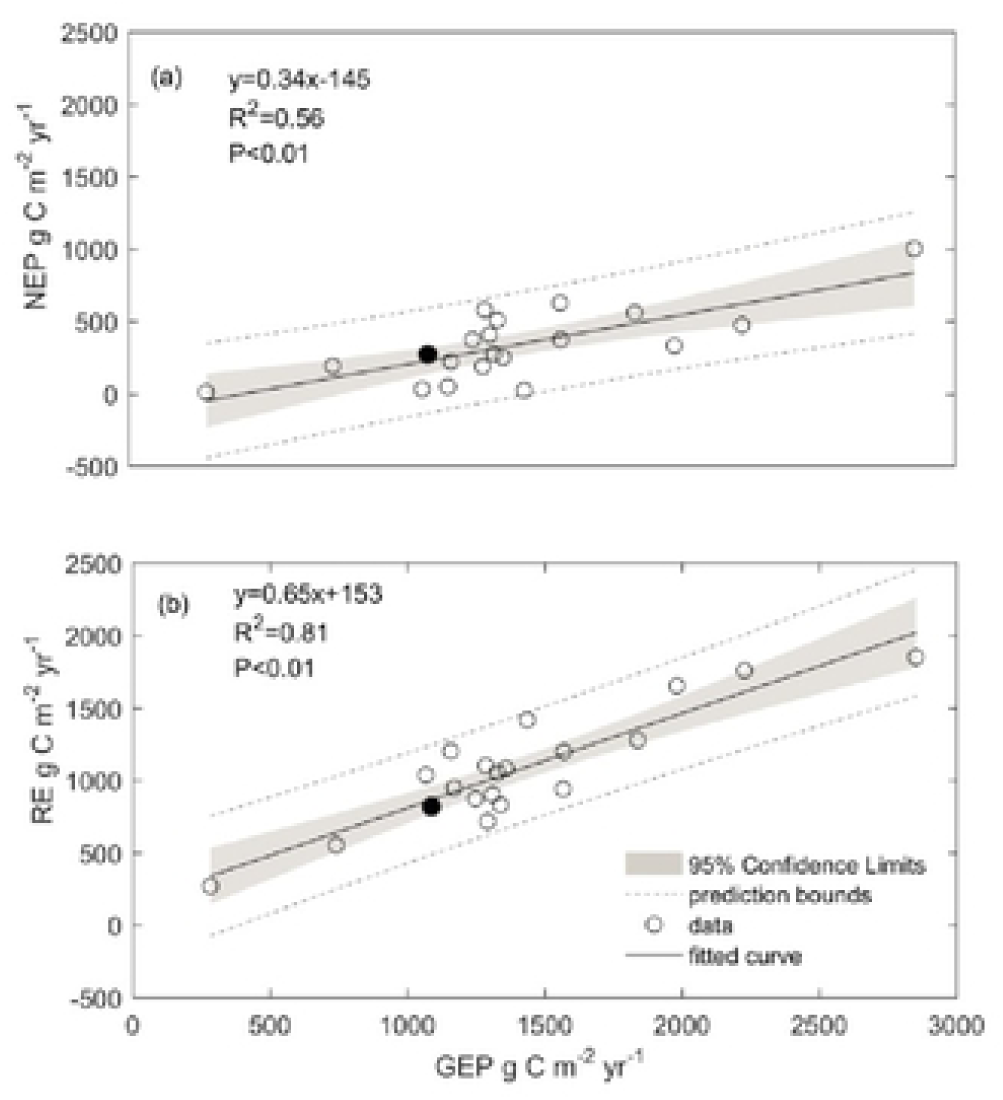
Dependence of annual net ecosystem production (NEP) (a) and ecosystem respiration (RE) (b) on annual gross ecosystem productivity (GEP) at multiple-site scale of 20 farmland sites from the literatures. Dataset listed in Tables S6, and black pot indicates the result of this study.

The IAV of GEP was controlled by temperature at the regional scale and driven by moisture, LAI and residues as well as temperature at the single site scale [4, 53]. Moreover, GEP was more sensitive to drought than RE in most ecosystems [15]. Both GEP and RE were restrained by water stress in drought years and the limitation of GEP was more than RE [20, 54]. Overall, the influences of climatic and biotic factors on GEP might vary with the changes of time and space scales. In this study, GEP was directly controlled by SWC and indirectly effected by LAI, VPD and PPT (Fig. 3).

The IAV of temperature can effect RE both at the regional and single site scale [6, 13, 55]. Moreover, RE was more sensitive to temperature than GEP, and soil moisture, LAI and biomass were also the drivers of RE [15, 54, 56]. The RE increased with the increasing of temperature at the regional scale according to 20 study sites from the published literature on different farmlands (p<0.05, Table S6). The variability caused by the IAV of temperature in RE was more than that in GEP [54, 56]. For example, Chen et al. [54] pointed out that both GEP and RE were enhanced by the increasing Ta with sufficient water supply, and the increasing of RE was more than that of GEP in the Pacific Northwest Douglas-fir forest. But the decline of soil moisture might result in weaker sensitivity of RE dependence on soil temperature [20]. Wilkinson et al. [13] also said that the responsiveness of RE to air temperature was strongly correlated to the summer soil moisture. Guo et al. [50] found that RE was primarily driven by soil water conditions in the maize farmland with a precipitation of 164 mm. In this study, RE was directly controlled by Ts and indirectly effected by LAI, VPD and PPT (Fig. 3).

### 4.2 Partitioning NEP into the integration of peak value and duration

The NEP is defined as the integration of the carbon uptake or release peak and the corresponding duration [1, 14-16]. The Previous studies suggest that climatic and biotic drivers may ultimately cause the IAV of NEP by regulating phenological and physiological indicators [4, 9, 15]. Therefore, the IAV of NEP can be explained by indicators related to physiological (described by the carbon uptake or release peak) and phenological (described by the corresponding duration) processes of the plant [53]. The framework provides a new perspective on the explaining the causes of the IAV of NEP [8, 14, 15, 23]. Because some climatic and biotic factors had the compensatory effects on the integration of peak value and duration, the negligible impacts on annual NEP and conflicting results in previous studies occurred [57, 58]. Meanwhile, carbon realease period was always ignored in previous studies [18, 28, 48, 59].

Typically, the IAV of NEP was positively correlated with annual anomalies of NEP_max_ and CUP, and annual anomalies of NEP_max_ had the large contributions compared to the other parameters from the framework of carbon uptake or release peak and the corresponding duration [1, 8, 14-16, 23]. Based on the 24 sites including broadleaf and evergreen forests and grasslands, it found that annual anomalies of NEP were dominant positively correlated with NEP_max_ and CUP [14], respectively. In this study, annual anomalies of NEP had a significant and positive correlation with NEP_max_ and CUP (Table 3), respectively. Zhou et al. [8] indicated that non-forest ecosystems generally had more obvious changes of an uptake peak and corresponding duration than forest ecosystesms because of poor self-regulation abilities under the environmental stress. Based on the 66 EC sites, the contribution of the IAV of NEP_max_ for NEP was 48% and that of CUP was 25%, and it indicated that the IAV of NEP was predominately determined by NEP_max_ [16]. In this study, NEP_max_ and CUP attributed to 84.1% and 9.7% of annual anomalies of NEP, respectively (Table 4). Zscheischler et al. [60] showed that the occurrence numbers of high values in observed daily ecosystem fluxes were strongly correlated with their annual sums, while the influence of phenological transitions was less important based on eight forest sites of the AmeriFlux network. Xia et al. [23] suggested that comprehensive controls of the uptake peak and corresponding duration on annual carbon uptake variability were strong in temperate and boreal ecosystems because these ecosystems only had one uptake peak with obvious seasonal dynamics, and were weak in tropical and Mediterranean climates.

Annual anomalies of NEP_max_ were mainly controlled by the moisture and temperature. According to the study of Fu et al. [14] annual anomalies of summer precipitation had a negative correlation with NEP_max_ in the broadleaf forest and a positive correlation in the grasslands. Nara et al. [51] stated that NEP_max_ during the wet years was higher than that during the drier years in the northern reed canary grass environment. Montagnani et al. [61] showed that the apple orchard ecosystem had the highest NEP_max_ during the wet year in northern Italy. Vendrame et al. [62] found that NEP_max_ was reduced by elevated air temperature and declined soil moisture in the temperate-climate vineyard. Buermann et al. [57] and Wolf et al. [58] indicated that warming at the beginning of the year led to earlier springs and reduced NEP_max_, because the increasing of evapotranspiration during the warming spring might cause the shortage of soil water in summer and the plant growth limitation. In this study, annual anomalies of summer VPD were negatively correlated with NEP_max_ (Fig 4).

Annual anomalies of CUP can be affected by temperature, precipitation, radiation, and LAI. The CUP were composed of BDOY and EDOY. The factors affecting on CRP were consistent with CUP. Fu et al. [14] indicated that BDOY had a negative correlation with spring temperature in the deciduous broadleaf and evergreen forests. The advancement of the growing season in response to a warming spring was well documented in previous studies, and it benefited the advance of BDOY [63, 64]. Agricultural ecosystems were also affected by the management practices, such as planting date compared to the natural ecosystem [65]. Knox et al. [6] showed that exogenous factors, such as the wetness of planting, could modulate crop growth, and thus suitable soil moisure and temperature were the important indicators for ensuring the best time of plant, and BDOY might be delayed with the retardation of sowing date. Fu et al. [14] showed that EDOY was positively correlated with autumn precipitation in grasslands, the autumn temperature in broadleaf forests, and autumn radiation in evergreen forests. In addition, adequate precipation, temperature, and a larger LAI can delay crop senescence in autumn, and it might lead to a delayed EDOY [35, 66]. In this study, BDOY had a significant increasing trend while EDOY were relatively stable, and it led to a significant decreasing tend in CUP (Table 2). Annual anomalies of the BDOY were negatively correlated with spring effective accumulative temperature and precipitation, and the EDOY was positively correlated with autumn precipitation and LAI (Fig. 4).

## 5. Conclusions

Climatic and biotic controls of the IAV of NEP were investigated based on an EC dataset of rain-fed spring maize from 2005–2018 in northeast China. Our results showed that the maize ecosystem acted as a carbon sink and the annual NEP was 270±115 g C m^−2^yr ^−1^. Annual values and anomalies of NEP were positively correlated with PPT, and annual anomalies were negatively correlated with PAR and VPD. Annual values and anomalies of NEP were positively correlated with GEP but not with RE. Annual values and anomalies of NEP were positively correlated with that of NEP_max_, and annual anomalies were positively correlated with that of CUP and negatively with that of CRP at the beginning of the year.

The SEM results showed that annual anomalies of NEP were dominantly and positively controlled by SWC through GEP by soil temperature through RE. Annual anomalies of NEP and the NEP of CUP were dominantly controlled by summer VPD through NEP_max_ and adjusted by spring precipitation and the effective accumulative temperature through BDOY affecting CUP, and by autumn precipitation and LAI through EDOY affecting CUP. Overall, residues produced in the previous year might decrease RE in the early part of the year because soil temperature was prevented from rising.

## Acknowledgments

This work was supported by the National Key Research and Development Program of China (2017YFC0503904) and the National Natural Science Foundation of China (41671257).

